# MYCN-induced nucleolar stress drives an early senescence-like transcriptional program in hTERT-immortalized RPE cells

**DOI:** 10.1101/2021.01.22.427454

**Authors:** Sofia Zanotti, Suzanne Vanhauwaert, Christophe Van Neste, Volodimir Olexiouk, Jolien Van Laere, Marlies Verschuuren, Liselot M. Mus, Kaat Durinck, Laurentijn Tilleman, Dieter Deforce, Filip Van Nieuwerburgh, Michael D. Hogarty, Bieke Decaesteker, Winnok H. De Vos, Frank Speleman

## Abstract

*MYCN* is an oncogenic driver in neural crest-derived neuroblastoma and medulloblastoma. To better understand the early effects of MYCN activation in a neural-crest lineage context, we profiled the transcriptome of immortalized human retina pigment epithelial cells with inducible MYCN activation. Gene signatures associated with elevated MYC/MYCN activity were induced after 24 h of MYCN activation, which attenuated but sustained at later time points. Unexpectedly, MYCN activation was accompanied by reduced cell growth. Gene set enrichment analysis revealed a senescence-like signature with strong induction of p53 and p21 but in the absence of canonical hallmarks of senescence such as beta-galactosidase positivity, suggesting incomplete cell fate commitment. When scrutinizing the putative drivers of this growth attenuation, differential gene expression analysis identified several regulators of nucleolar stress. This process was also reflected by phenotypic correlates such as cytoplasmic granule accrual and nucleolar coalescence. Hence, we propose that the induction of MYCN congests the translational machinery, causing nucleolar stress and driving cells into a transient pre-senescent state. Our findings shed new light on the early events induced by MYCN activation and may help unravelling which factors are required for cells to tolerate unscheduled MYCN overexpression during early malignant transformation.

**Highlights:**

- Activation of MYCN attenuates proliferation in RPE1 cells
- Growth arrest is associated with an early senescence-like transcriptional signature
- Transcriptional and phenotypic evidence of nucleolar stress
- *CDKN1A* upregulation in G2 phase primes cells for faltering in subsequent G1

## 1. Introduction

MYCN belongs to the MYC protein family of bHLH transcription factors, also including MYC and MYCL. While MYC is commonly activated in many cancers, MYCN is also increasingly recognized as important oncogenic driver in a growing number of cancer entities. MYCN was first discovered in neuroblastoma and later also found to be overexpressed in retinoblastoma, brain tumors, leukemia, neuro-endocrine prostate cancers and pancreatic cancer (Rickman, D. S. et al., Rickman, D. S.018). MYCN has been extensively studied in neuroblastoma where it marks a subset of high-risk children with poor prognosis despite intensive multi-modal therapy. While mutational burden in these tumors is low, MYCN amplified neuroblastomas and medulloblastoma often exhibit large copy number alterations, most notably 17q gain (Brady, S. W. et al., 2020; Hovestadt, V. et al., 2019), indicating that additional gene dosage effects are required to support tumor initiation and/or progression. Given their structural properties, the MYC proteins have been considered as undruggable, but new insights into their function and protein interactions are providing exciting insights towards novel strategies to target these oncogenes. Despite decades of intensive research, the mechanism through which MYC(N) proteins drive cancer formation remain poorly understood. Most of the earlier work focussed on the identification of directly regulated target genes assuming a classical transcription factor role. More recently, evidence was obtained that MYC(N) influences global transcriptional changes through a so-called “amplifier” mode of action. Finally, based on protein-interactions and further in-depth biochemical studies by the Eilers team (Rickman, D. S. et al., 2018), MYC(N) oncogenicity was attributed to its role in enhancing transcription-stress resilience in tumour cells. This includes roles in alleviating replication fork stalling resulting from nucleotide depletion and POL II stalling, R-loop formation and replication-transcription conflicts (Baluapuri, A. et al., 2020). Despite these increasing insights, our current understanding of the early cellular responses that allow cells to cope with the consequence of MYCN overexpression in untransformed cells are limited. Hence, we explored the early changes in the dynamic transcriptional landscape of the hTERT-immortalized tamoxifen-inducible RPE1–MYCN-ER human cell model.

In contrast with the known growth promoting effect of MYCN, we observed attenuated cell growth accompanied with nucleolar coalescence, appearance of cytoplasmic granules and increased p21 and p53 protein levels. Yet, senescence hallmarks such as beta-galactosidase positivity, DNA damage or heterochromatin foci formation were absent. We further found that MYCN-induced growth deceleration is accompanied with a transcriptional signature that resembles early senescence induction thus offering a platform for further mechanistic studies to explore cellular processes that protect against early MYCN-induced cellular transformation.

## 2. Materials and Methods

### 2.1 Cell culture

The hTERT-immortalized MYCN retinal pigmented epithelial cell line (RPE1–MYCN-ER, kindly provided by Michael Hogarty, Children’s Hospital of Philadelphia, Philadelphia, PA) carries a MYCN:ER expression construct that results in its constitutive expression as described by Maris *et al*. (Wang, Q. et al., 2003). When 4-OHT is present in the culture media, MycN:ER translocates to the nucleus where it is transcriptionally active. The wilde-type (WT) hTERT-immortalized retinal pigment epithelium (hTERT-RPE1, Clontech, Palo Alto, CA) and RPE1–MYCN-ER were cultured in DMEM/F-12 HEPES, supplemented with 10% fetal bovine serum (FBS), 1% penicillin/streptomycin, 1% L-glutamin and 0,01 mg/ml Hygromycin B according to standard procedures. Moreover, 1 µg/ml puromycin was added to the RPE1–MYCN-ER medium composition as a selection marker for the 4-OHT inducible construct. Cell culture products were obtained from Life Technologies and mycoplasma testing was done on a two-monthly basis.

### 2.2 Cell seeding and treatment

To evaluate phenotypic and protein changes, RPE1–MYCN-ER cells were plated in 96 wells overnight in three biological replicates and treated with or without 400 nM 4-OHT (H6278; Sigma-Aldrich) for 24 h (4000 cells per well), 48 h (2000 cells per well) and 72 h (1000 cells per well). For the detection of stress granules and P-bodies, a treatment with 0.5 mM NaAsO_2_ (#S7400; Sigma-Aldrich) for 1 h served as positive control. For DNA damage detection, the positive control consisted of a treatment with 1 μM 3-AP (#2251; Tocris) for 72 h.

### 2.3 Focus formation (NIH-3T3) assay

To quantify the impact of the oncogene MYCN on the proliferation rate over time and several passages, a NIH-3T3-assay was applied as previously described by Todaro et al. (Todaro and Green, 1963). In brief, cells with an inducible MYCN OFF/ON system, respectively either maintained in control conditions (EtOH) or induced MYCN (4-OHT), were plated in T25 flasks at a density of 0,15 million cells. The cumulative increase in cell number was calculated according to the formula log(Nf/Ni)/log2, where Ni and Nf are the initial and final numbers of cells plated and counted after three or four days respectively.

### 2.4 Nuclear and cytoplasmic fractionation

The protocol was conducted as described earlier (Yue, M. et al., 2017). In brief, cells were grown as monolayers in T75 flasks, scraped from culture flasks incubated in 1.5 mL of Harvest buffer (10 mM Hepes (pH 7.9), 50 mM Nacl, 0.5 M Sucrose, 0.1 mM EDTA, 0.1% Triton x-100 with freshly added protease and phosphatase inhibitors) at 4°C for 15 min and collected in 1,5 ml micro-centrifuge tubes. After centrifugation (2000xg for 10 min), supernatant (cytoplasmic fraction) was collected from each sample and subjected to 13000xg for another 10 min in addition to cell pellets to remove any nuclear contamination and transferred to a new tube. The pellets were washed three times (3 times 2000Xg for 3 min), with washing buffer (10 mM Hepes (pH 7.9), 10 mM KCl, 0.1 mM EDTA, 0.1 mM EGTA and freshly added protease and phosphatase inhibitors). Both the pellet and supernatant were boiled separately in Laemmli sample buffer and β-mercaptoethanol. Western blot was performed using PVDF membranes incubated with anti-vinculin (Sigma-Aldrich; V9131, 1:10,000 dilution) antibody as cytoplasmic loading control, anti-HDAC (Santa-Cruz; sc-7872, 1:500 dilution) as nuclear loading control and anti-MYCN as a target protein (Santa-Cruz; sc-53993, 1:1000 dilution) after blocking with 5% non-fat milk in 1% tween-20 in TBST. Membranes were washed with TBST followed by incubation with HRP-conjugated anti-rabbit (Cell Signalling Technologies; #7074S, 1:4000 dilution) or anti-mouse secondary antibody (Cell Signalling Technologies; #7076S, 1:3000 dilution). After washing with TBST, imaging was performed using SuperSignal West DURA Extended Duration Substrate (Thermo Scientific; 34076×4) and the Amersham Imager 680 (GE Healthcare).

### 2.5 Real-time quantitative PCR

Total RNA was extracted using mRNeasy kit (Qiagen) according to the manufacturer’s instructions and concentration was determined with the Nanodrop (Thermo Scientific). cDNA synthesis was performed using the iScript Advanced cDNA synthesis kit from BioRad. PCR mix contained 5□ng of cDNA, 2.5□µl SsoAdvanced SYBR qPCR supermix (Bio-Rad) and 0.25□µl forward and reverse primer (to a final concentration of 250□nM, IDT) and was analysed on the LC-480 device (Roche) for RT-qPCR cycling. Expression levels were normalized using expression data of 3 stable reference genes (18s, HMBS, SDHA) and analysed using qBasePlus software (http://www.biogazelle.com). Relative abundance of *ODC1* transcripts (forward: GATGACTTTTGATAGTGAAGTTGAGTTGA; reverse: GGCACCGAATTTCACACTGA), *CDKNA1* transcripts (forward: CCTCATCCCGTGTTCTCCTTT; reverse: GTACCACCCAGCGGACAAGT), *RRM2* transcripts (forward: AGGACATTCAGCACTGGGAA; reverse: CCATAGAAACAGCGGGCTTC were measured relative to *18s* (forward: TTCGGAACTGAGGCCATGAT; reverse: TTTCGCTCTGGTCCGTCTTG), *HMBS* (forward: GGCAATGCGGCTGCAA; reverse: GGGTACCCACGCGAATCAC) and *SDHA* (forward: TGGGAACAAGAGGGCATCTG; reverse: CCACCACTGCATCAAATTCATG) reference transcripts.

### 2.6 Caspase-Glo 3/7 assay reagent

Caspase-3/7 activity was detected using a luminometric assay kit (Caspase-Glo 3/7; Promega) according to the manufacturer’s instructions. The reagent provides a proluminescent caspase-3/7 substrate, which contains the tetrapeptide sequence DEVD, in combination with luciferase and a cell-lysing agent. The addition of the Caspase-Glo 3/7 reagent directly to the assay well results in cell lysis, followed by caspase cleavage of the DEVD substrate, and the generation of luminescence. Luminescence detection was performed by means of a microplate reader (GloMax; Promega).

### 2.7 Immunofluorescence staining

RPE1–MYCN-ER cells were grown in black 96-well µClear plates (# 655090, Greiner Bio-One) and fixed with 4% paraformaldehyde for 20 minutes at room temperature followed by washing (3x, 5 minutes) with PBS. Subsequently, cells were permeabilized with 0.3% Triton X-100 (8 minutes) and washed (2x, 5 minutes), after which they were blocked with 50% FBS for 30 minutes and incubated with primary antibody diluted in 50% FBS for 60 minutes. After minimally 3 PBS wash steps, plates were incubated with secondary antibody diluted in 50% FBS for 60 minutes, washed again, and counterstained with DAPI (5 μg/ml) for 20 min. Primary antibodies were directed against p21 (F-5) (SC-6246, Santa Cruz Biotechnology Inc., 1:100 dilution), p53 (7F5) (#2527T, Cell Signalling Technology, 1:1600 dilution), Lamin B1 (ab16048, Abcam, 1:1000 dilution), Ki-67 (M7240) (M724029-2, Agilent, 1:400 dilution), γH2AX (phospho-S139) (ab2893, Abcam, 1:1000 dilution), RPA32/RPA2 (9H8) (ab2175, Abcam, 1:300 dilution), Fibrillarin (H-140) (SC-25397, Santa Cruz Biotechnology Inc., 1:50 dilution), DDX6 (200–192, Novus, 1:300 dilution) and G3BP1 (H00010146-M01, Abnova, 1:200 dilution). As secondary antibodies goat anti-rabbit IgG (H+L) Alexa Fluor 488 (A11034, Thermo Fisher, 1:400 dilution) and goat anti-mouse IgG (H+L) CY3 (A10521, Thermo Fisher, 1:400 dilution) were used. For cytoplasmic staining HCS CellMask™ Red stain (H32712, Thermo Fisher) was added according to the manufacturer’s instructions. After additional washing, the plates were maintained in 0.1% NaN_3_/PBS and stored at 4 °C for microscopy.

### 2.8 Lysosome staining

Live cells were stained with 5 μg/ml of the nuclear dye Hoechst 33342 (# H3570, Invitrogen) in medium at 37°C for 5 min. After the plate was washed twice with PBS, 100 μl/well 50 nM Lysotracker-red DND-99 dye (#L7528, Invitrogen) was added in medium at 37°C for 60 min. Thereafter, medium was replenished and prepared for imaging.

### 2.9 EdU incorporation assay

EdU (5-ethynyl-2′-deoxyuridine), supplied with Click-iT EdU Alexa Fluor 647 Imaging Kit (#C10340, Thermo-Fisher Scientific, Waltham, MA, USA), was diluted in DMSO to a final concentration of 10 mM and kept at -20°C. All steps of the Click-iT™ reaction were performed at room temperature. Typically, EdU was added to parallel cultures growing exponentially in 96 well plates to final concentration of 20 μM for 30 minutes to label the S-phase cells before fixation with 4% paraformaldehyde. Additional antibodies were added as previously described in the immunofluorescence protocol. During the secondary antibody incubation step, the Click-iT™ kit azide and buffer additive were removed from -20°C storage and allowed to thaw in a light-protected box. The buffer additive is stored as a 10X solution; once it is thawed, it must be diluted with dH2O to make a working 1X solution. During the third PBS rinse, the reaction cocktail is made, adding the azide last. The reaction cocktail must be used within 15 minutes of creation. The amount of reaction cocktail needed was determined according to the Click-iT EdU Alexa Fluor 647 Imaging Kit as per manufacturer’s instructions. Fixed cells were then incubated in the EdU cocktail for 30 minutes and rinsed three times in PBS for 5 minutes per rinse. The azide used was coupled to an Alexa Fluor® 647 (red) fluorophore. Upon completion of the EdU Click-iT™ reaction, DAPI (5 μg/ml) was added for 20 min to the cells and after additional washing the plates were maintained in 0.1% NaN3/PBS and stored at 4 °C for microscopy.

### 2.10 Microscopy

Image acquisition of fluorescently labelled cells (live or fixed) was performed using a Nikon Ti widefield fluorescence microscope, equipped with an Andor DU-885 X-266 camera. Per well, 16 fields were acquired in 4 channels (405, 488, 561 and 640□nm excitation) recorded using a 40x dry (NA = 0.95) objective. Live cells were imaged in DMEM/F-12 HEPES medium under environmental control (37 °C, 5% CO2).

### 2.11 Image processing and data analysis

Image processing was performed in FIJI (http://fiji.sc), a packaged version of ImageJ freeware (W.S. Rasband, USA. National Institutes of Health, Bethesda, Maryland, USA, http://rsb.info.nih.gov/ij/, 1997–2014). Quantification of nuclear and spot signal intensities of (immuno-)stained cell cultures was done using a script specifically written for automated cell-based analysis (CellBlocks.ijm) (De Vos, W. H. et al., 2010). In brief, the image analysis pipeline relies on the segmentation of nuclei, cells and intracellular spots and subsequent feature (number, intensity, shape…) extraction for all regions of interest. Downstream data analysis was performed in R Studio and graphs were generated by Graphpad Prism (version 8). To identify numbers (and percentages) of marker-positive and -negative cell nuclei, a manual threshold was set on the intensity values of the marker of interest. For cell cycle staging, we plotted the integrated nuclear signal intensity of the DAPI channel (Roukos, V. et al., 2015). The results shown are averages (with standard deviation interval, tested by unpaired t-test) of a minimum of three independent experiments. Significance levels were indicated as follows: p <0.05 (*), p <0.01 (**), and p <0.001 (***).

### 2.12 Bulk RNA-sequencing

To evaluate transcriptomic changes RPE1–MYCN-ER and WT hTERT-RPE1 cells were plated in T75 flasks overnight in four biological replicates and treated with or without 400 nM 4-OHT (H6278; Sigma-Aldrich) for 24 h, 48 h and 72 h. Controls were supplied with the same amount of EtOH as was present in the treated cells and was always ≤0.5% of the total volume. Total RNA was extracted using mRNeasy kit (Qiagen) according to the manufacturer’s instructions. After RNA extraction, the concentration and quality of the total extracted RNA were checked by using the ‘Quant-it ribogreen RNA assay’ (Life Technologies, Grand Island, NY, USA) and the RNA 6000 Nano chip (Agilent Technologies, Santa Clara, CA, USA), respectively. Subsequently, 400 ng of RNA was used to perform an Illumina sequencing library preparation using the QuantSeq 3’ mRNA-seq Library Prep Kit (Lexogen, Vienna, Austria) according to the manufacturer’s protocol. During library preparation, 14 PCR cycles were used. Libraries were quantified by qPCR, according to Illumina’s protocol (Sequencing Library qPCR Quantification protocol guide, version February 2011). A High Sensitivity DNA chip (Agilent Technologies, Santa Clara, CA, US) was used to control the library’s size distribution and quality. Sequencing was performed on a high throughput Illumina NextSeq 500 flow cell, generating 75 bp single reads. Per sample, on average, 7.5 × 10^6^ ± 1.6 × 10^6^ reads were generated. First, these reads were trimmed using cutadapt (Martin, M. et al., 2011) version 1.16 to remove the ‘QuantSEQ FWD’ adaptor sequence. The trimmed reads were mapped against the Homo sapiens GRCh38.89 reference genome using STAR (Dobin, A. et al., 2013) version 2.6.0c. The RSEM (Li, B. et al., 2011) software, version 1.3.1, was used to generate the count tables. To explore if the samples from different groups clustered together and to detect outlier samples, Principal Component Analyses (PCAs) on rlog transformed counts were performed using R, statistical computing software. Genes were only retained if they were expressed at counts per million (cpm) above one in at least four samples. Counts were normalized with the TMM method (R-package edgeR), followed by voom transformation and differential expression analysis using limma (R-package limma). Gene set enrichment analysis was performed on the ranked genes according to differential expression statistical value (t) and corresponding heatmaps were made using a custom R script. Signature scores were conducted using a rank-scoring algorithm (Fredlund, E. et al., 2008). For the time-series analysis, all genes were clustered according to their average counts across replicates. Counts were quantile normalized across genes and across samples. Clusters were made with k-means method from the Python sklearn package (v0.22.2). To determine the ideal number of clusters, the elbow method was used (Ketchen, Jr. D. and Shook, C., 1996).

### 2.13 Single cell RNA-sequencing

Cells were harvested after treatment and washed two times with PBS with 0.04% BSA. The viability and the number of cells were measured with the Countess II (Invitrogen, Carlsbad, CA, USA). Samples were prepared for loading on the 10x Genomics Chromium microfluidics controller, as described in the Chromium Single Cell 3’ protocol (10x Genomics, Pleasanton, CA, USA). Per sample, 5000 cells were loaded to end up with 2500 captured cells. During cDNA amplification, 12 PCR cycles were used. To add an index to the samples, another 12 PCR cycles were used. To check the final quality of the generated libraries, the samples were loaded on a Bioanalyzer High sensitivity chip (Agilent Technologies, Santa Clara, CA, US), and the quantity was checked with qPCR according to the Illumina protocol ‘Sequencing Library qPCR Quantification protocol guide’, version February 2011. Samples were pooled equimolarly and sequenced on a Hiseq4000 lane (Illumina, San Diego, CA, USA), generating paired-end reads of 150 base pairs. Afterward, the data were trimmed to paired-end reads of 28 and 91 base pairs. Single cell analysis was performed with the Python scanpy package (v1.5.1). Cells that reported less than 1000 genes were filtered, so were genes that were reported in only 1 or 2 cells. Cells were allowed to have up to 12% mitochondrial reads. For determining the differential expression of single cell genes, the Python package diffxpy (v0.7.4) was used. To calculate the bulk RPE1-MYCN-ER signatures in the single cell uniform manifold approximation and projection (UMAP) spaces, only genes were considered with an adjusted p-value < 0.01. Single cell library sizes were adjusted to 10000 and transformed by natural logarithm after addition of 1. UMAPs with 3 components were produced by taking 10 neighbours in a 25 component PCA space. Cell phase was determined with the S and G2M gene sets used in (Pont, F. et al., 2019). In figures with all treatments together, cell phase was determined across all data, in figures were only one treatment is displayed, cell phase was determined for each cell only within that treatment data. Gene senescence signature scores were calculated as the number of genes that had expression in the cells. For the bulk RPE1-MYCN-ER signatures in the single cell, the average rank of the genes in the set was calculated. Full scripts can be found at https://github.com/dicaso/apetype/tree/master/examples/RPEMYCN.

### 2.14 Quantification and statistical analysis

Statistical analyses were performed in R Studio and statistical significance of differences between conditions for the immunofluorescence and caspase-glo experiments were determined by unpaired Student’s t-test using Prism (version 8). The ANOVA (analysis of variance) test followed by a post-hoc Tukey’s test for multiple comparisons was used to determine differences in gene expression and signatures scores between 4 different groups with 3 (RT-qPCR) or 4 (RNA-Seq) biological replicates per condition. For RT-qPCR experiments, reference genes were excluded if GeNorm M value was greater than five and/or Coefficient of Variation greater than two, according to the qBase+ software. For all experiments, at least three reference genes were used for the normalization according to good RT-qPCR practice. All error bars represent the SD or 95% CI of the three or four biological replicates. For statistical testing of the focus formation (NIH-3T3) assay, cumulative growth values were modelled using a linear model with time and treatment as fixed factors and an interaction term for both (Python package statsmodels). This model explained 99% of the observed variance and performed better than a restricted model omitting the interaction term for time and treatment (LR test P-value < 0.01, interaction term P-value < 0.01 in the more complex model). The details of quantification and statistical methods used are mentioned in the figure legends.

### 2.15 Data availability

The bulk and single-cell RNA-sequencing datasets generated during this study were deposited in the ArrayExpress database at EMBL-EBI EBI (www.ebi.ac.uk/arrayexpress) with accession numbers E-MTAB-9667 and E-MTAB-9689, respectively.

## 3. Results

### 3.1 MYC/MYCN gene signature analysis validates MYCN activation in immortalized RPE1 cells

To study the early effects of MYCN activation, we used 4-OH tamoxifen (4-OHT) inducible *MYCN* gene regulation in telomerase immortalized RPE1 cells (further referred to as RPE1-MYCN-ER) (Wang, Q. et al., 2003). In this model, chaperonins persistently sequester the chimeric MYCN-ER protein in the cytosol. The oestrogen receptor ligand 4-OHT will displace the chaperonins, causing MYCN-ER to translocate to the nucleus where it will exert its transcriptional activity. Western blotting confirmed the expected decrease in the cytoplasmic fraction of MYCN-ER protein (Fig. 1A), and concomitant increase in nuclear levels 24h after the addition of 4-OHT to RPE1-MYCN-ER cells (MYCN-ON) as compared to mock-treated RPE1-MYCN-ER (MYCN-OFF) and wild type RPE1 (WT) cells (Fig. 1B). First, we evaluated the functionality of MYCN after nuclear translocation in RPE1 cells by measuring the transcriptional impact. To this end, bulk RNA sequencing was performed at three sequential time points post induction (p.i.) (24 h, 48 h, 72 h) for MYCN-ON and 4-OHT-treated WT cells, as well as MYCN-OFF and mock-treated WT cells (0 h) (Supplementary Table 1). Principal component analysis (PCA) revealed a clear separation of the MYCN-ON cells from the other conditions along the first principal component (PC) (Fig. 1C). Within the MYCN-ON samples, there was also a pronounced separation of individual time points, predominantly dictated by the second PC. This demonstrates that incubation with 4-OHT induces a specific and time-dependent transcriptional shift in MYCN-ON cells (Fig. 1C).

**Fig 1.**
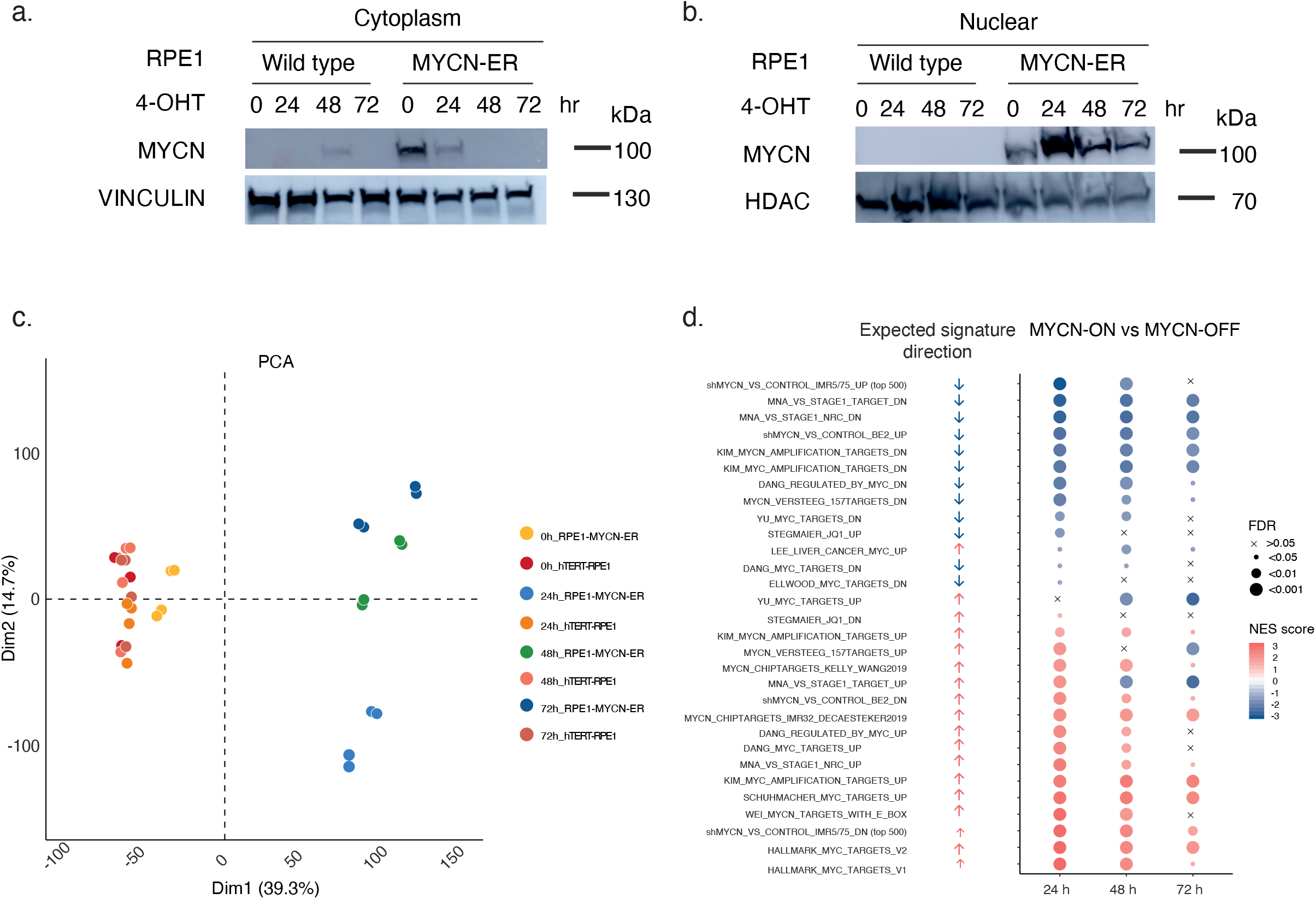
MYCN activation induces MYC/MYCN gene signatures with a specific and time-dependent transcriptional shift in MYCN-ON cells. Immunoblot of N-MYC nuclear (A) and cytoplasm (B) translocation in WT and RPE1-MYCN-ER cells upon 4-OHT treatment for 24, 48 and 72 h. Vinculin was used as a cytoplasmic marker and loading control (A) while HDAC1 was used as a nuclear marker and loading control (B). (C) Principal component analysis and clustering with all samples, before differential analysis. All genes with a cpm greater than 1 in at least 6 samples are used in the analysis. Counts are normalized using edgeR and the data are rlog transformed. (D) 30 public available bona fide MYCN/MYC targets gene sets enriched in MYCN-ON cells 24 h, 48 h and 72 h p.i. compared to MYCN-OFF cells. Arrows represent the theoretically expected regulation of each signature in a positive (red) or negative (blue) manner for RPE1-MYCN-ER cells after MYCN activation. FDR value is depicted by dot size, GSEA normalised enrichment score (NES) is depicted by the colour of the dot.

To assess the effect of MYCN activation on the RPE1 transcriptome, we performed gene set enrichment analysis (GSEA) using 30 publicly available *MYCN/MYC* target gene sets. This revealed the expected enrichment (FDR<0.01) of *MYCN/MYC* targets in MYCN-ON cells at 24 h and onwards while this was not visible for the MYCN-OFF cells (Fig. 1D). In support thereof, RNA-sequencing and qPCR indeed revealed an upregulation of several selected MYCN target genes (*e*.*g*., *ODC1, PTMA, WDR12, MDM2, NPM1, DKC1* (Oliynyk, G. et al., 2019; Westermann, F. et al., 2008) (Supplementary Fig. 1A-B). When evaluating MYCN activity (translocation and signature activation) over time, the highest activity was noted at the 24 h time point, followed by sustained but attenuated activity at the later 48 h and 72 h time points (Fig. 1B, Supplementary Fig. 1C). This may be the result of a negative feedback loop as previously described (Cole, K. A. et al., 2011), a notion supported by the lower *MYCN* transcript levels in MYCN-ON cells as compared to MYCN-OFF cells at all time points (Supplementary Fig. 1D).

### 3.2 MYCN activation induces a p53-p21 driven growth attenuation

We noticed that MYCN-ON cells did not reach the same level of confluency 72 h p.i. as compared to controls (Fig. 2A). This pointed to an attenuation in cell proliferation, which was confirmed with a focus formation assay (P-value<0.01) (Fig. 2B). Quantitative immunofluorescence microscopy revealed that MYCN-ON cell populations harboured consistently fewer replicating (EdU-positive) cells than controls and progressively fewer cycling (Ki-67-positive) cells, with time up to 72h after *MYCN* induction (Fig. 2C,D,E). Caspase-3 and -7 activity was not significantly increased at any of the measured time points in MYCN-ON cells, suggesting there was no induction of apoptosis (Supplementary Fig. 2). DAPI-based cell cycle staging showed an increased fraction of cells in G1 phase and a significant decrease in G2 phase in MYCN-ON cells starting at 48 h p.i. (Fig. 2F).

**Fig 2.**
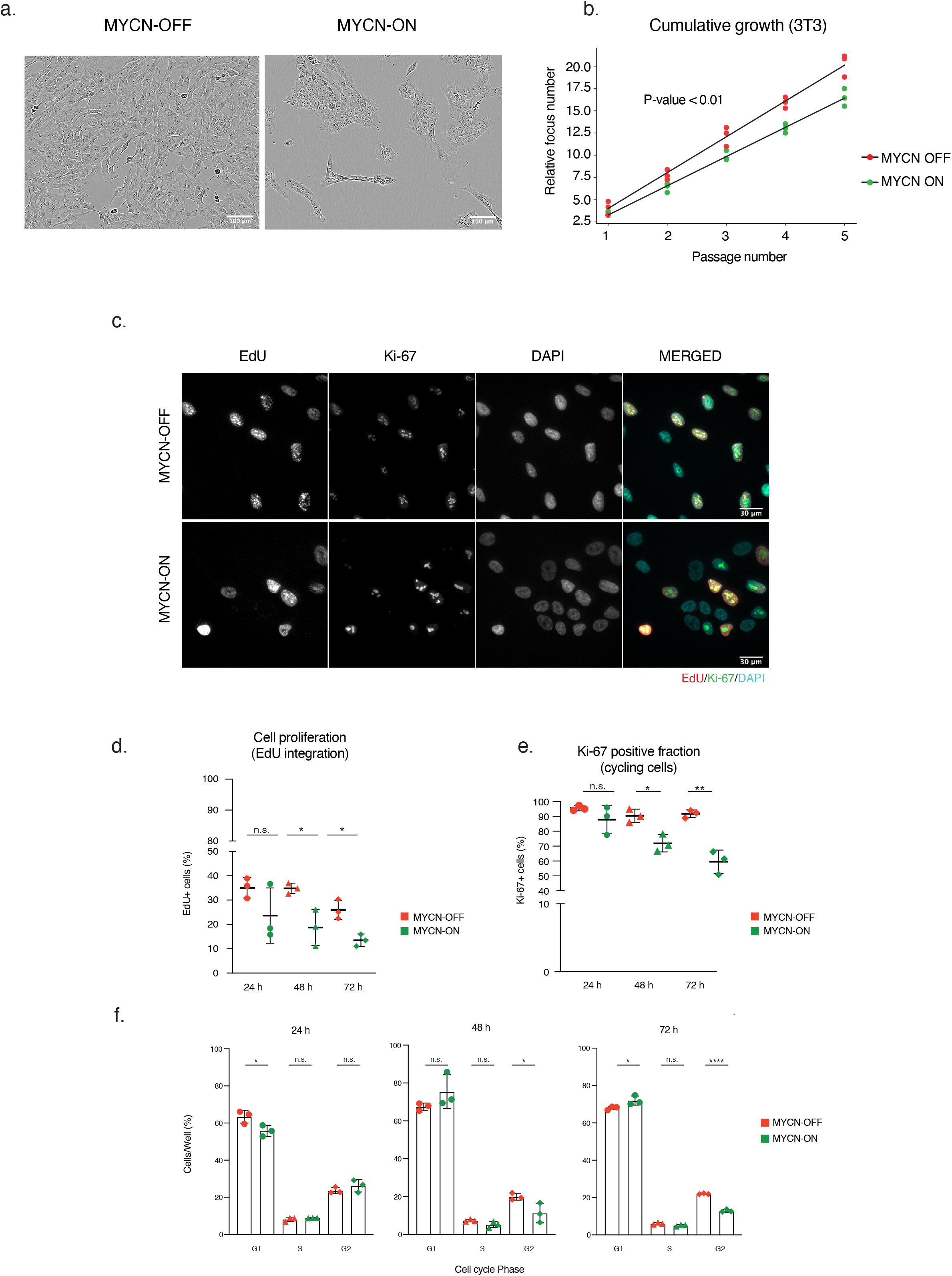
MYCN-ON cells undergo a slow-down in proliferation evidenced by reduced EdU incoporation and Ki-67 staining. (A) Phase contrast image of the corresponding cell line in both conditions 72 h p.i.. (B) focus-forming (NIH3T3) assay measuring the oncogenic potential of MYCN by calculating the cumulative increase in cell number. (C) Representative images of MYCN ON and OFF cells at 72 h p.i., after immunofluorescence staining for EdU (red), Ki-67 (green) and counterstaining with DAPI (cyan). (D) EdU incorporation immunofluorescence, expressed by the number of EdU positive nuclei relative to the total number of cells, in MYCN-ON and MYCN-OFF cells over time and (E) quantification of Ki-67, expressed by the number of Ki-67 positive nuclei relative to the total number of cells in MYCN-ON and MYCN-OFF cells over time. Error bars represent SD of the three biological replicates. (F) DAPI-based cell cycle staging upon MYCN activation. Statistical differences examined by unpaired Student’s t-test. Error bars represent SD of the three biological replicates. (n.s.= not significant, * = p-value < 0, 05; ** = p-value < 0, 01; *** = p-value < 0, 001). Scale bar□=□400□μm.

To gain insight into the underlying cause of growth attenuation, we analysed accompanying changes in the transcriptome of MYCN-ON cells. Using gene set enrichment analysis (GSEA), we found enrichment for predicted p53 promotor-binding sites in the upregulated genes while the downregulated genes were enriched for FOXM1- and E2F1-binding targets, 48 h and 72 h after MYCN induction (Fig. 3A-B, Supplementary Fig. 3A). Positive regulators of cell cycle transcription factors E2F1 and E2F2 were strongly downregulated after 24 h (Supplementary Fig. 3A). At all tested time points, transcriptional upregulation was found for Cyclin Dependent Kinase Inhibitor 1A (*CDKNA1*), encoding the p21 protein and a direct target gene of p53. Induction of p21 by p53 is implicated in cell cycle arrest and senescence (Georgakilas, A. G. et al., 2017; Rufini, A. et al., 2013) in accordance with the observed growth attenuation (Fig. 3C; Supplementary Fig. 3B). Quantitative immunofluorescence of MYCN-ON versus MYCN-OFF cells, revealed increased p53 and p21 protein (Fig. 3D), in the absence of notable *TP53* mRNA changes (Supplementary Fig. 3C). Deconvolution of nuclear marker intensities along the cell cycle revealed that p21 was much more abundant in G1-phase as compared to S or G2 phase (Supplementary Fig. 3D). This was already evident prior to MYCN induction (*i*.*e*., in MYCN-OFF), but became even more pronounced in MYCN-ON cells, suggesting this population to selectively enrich. In contrast, p53 protein levels increased in MYCN-ON cells independently of cell cycle status (Supplementary Fig. 3D-E).

**Fig 3.**
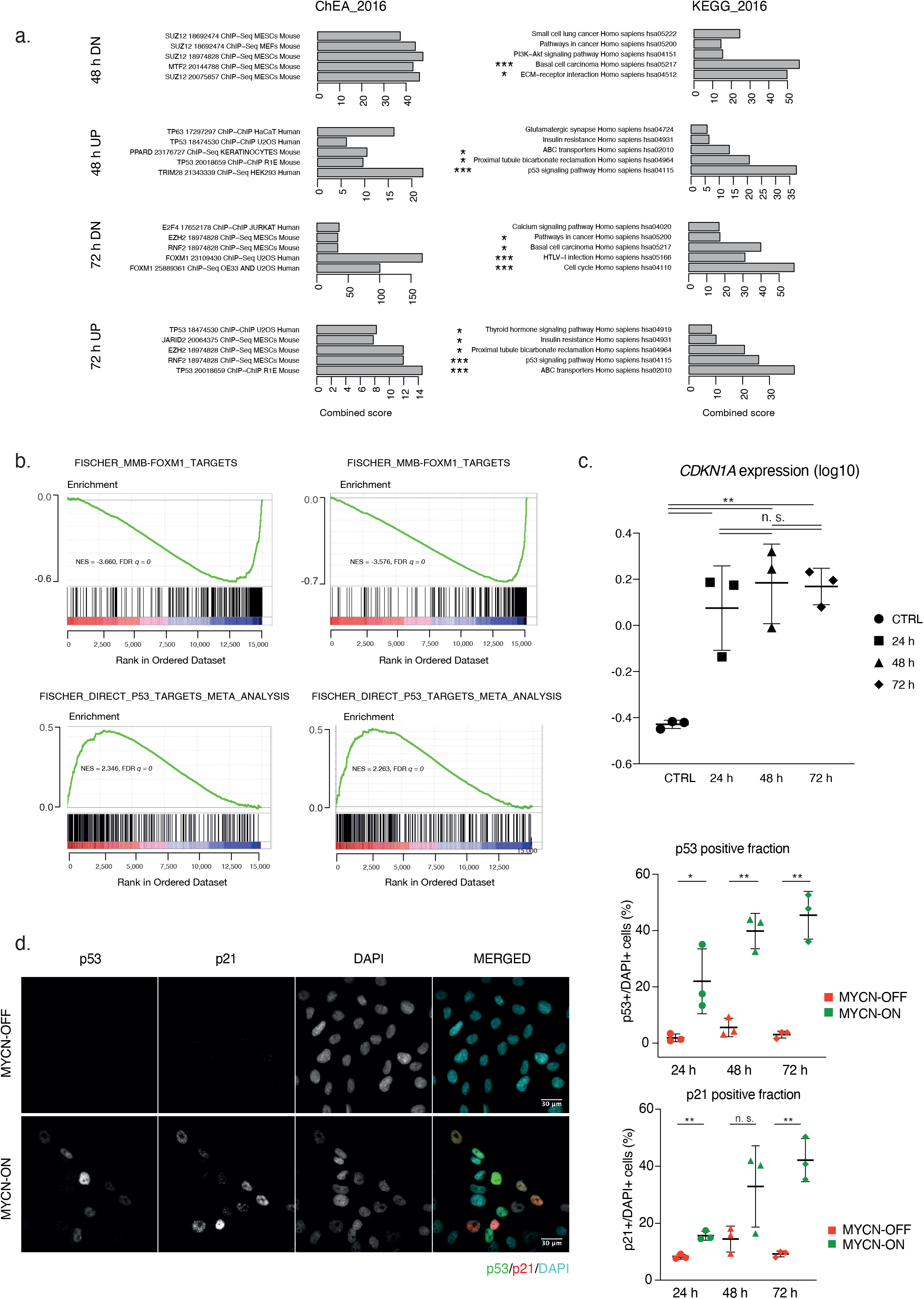
MYCN activation triggers a p53-p21 driven growth attenuation. (A) Enrichr (https://maayanlab.cloud/Enrichr/) analysis shows enrichment for predicted p53 promotor binding sites in the upregulated genes and enrichment for FOXM1 and E2F1 binding in the downregulated genes 48 h and 72 h p.i.. Combined Z-score is depicted based on the multiplication of log p value computed with Fisher exact test and the z-score which is the deviation from the expected rank by the Fisher exact test. The top 5 ranked gene sets based on combined z-score are depicted, which are all significant (p<0.05). Gene sets marked with an asterisk indicate significance upon multiple testing correction with *** =adj p.val < 0.001 and * adj p.val < 0.05. (B) FOXM1 (top) and p53 (bottom) targets related gene sets are enriched in MYCN-ON 48 h p.i. (left) and 72 h p.i. (right) compared to untreated MYCN-OFF. (C) log10 *CDKN1A* mRNA levels 24□h, 48 h, 72 h p.i.. Results are normalised to *18s, HMBS* and *SDHA* expression. (D) Representative images of MYCN-ON and OFF cells 72 h p.i., after immunofluorescence staining for p53 (green), p21 (red) and counterstaining with DAPI (cyan), with relative protein levels of p53 (top) and p21 (bottom) in MYCN ON versus OFF, as measured by immunofluorescence staining and quantitative image analysis. Statistical differences examined by unpaired Student’s t-test. Error bars represent SD of the three biological replicates. (n.s.= not significant, * = p-value < 0, 05; ** = p-value < 0, 01; *** = p-value < 0, 001). Scale bar□=□30□μm.

Taken together, MYCN activation induces p53-p21 transcript and protein levels along with upregulation of p53 target genes and attenuated expression of FOXM1 and E2F target genes.

### 3.4 Induction of a pre-senescent transcriptional program in MYCN-ON cells

The combination of a marked growth attenuation and upregulation of p53 and p21 pathways, could be indicative for the induction of MYCN-triggered (oncogene-induced) senescence. To investigate this, we queried our RNA-seq dataset for a variety of senescence-related signatures (Fig. 4A) and observed a strong enrichment as of the 48 h time point after MYCN induction for the cell cycle arrest and senescence associated gene sets (Fig. 4A). In this context, the enrichment for a YAP/TAZ gene signature (Rajbhandari, P. et al., 2018; Hiemer, S. E. et al., 2015) (Fig. 4A, 4B) is highly relevant given the reported downregulation of YAP proteins following replication-induced cellular senescence in IMR90 cells (Xie, Q. et al., 2013). YAP/TAZ signaling has also been shown to control *RRM2* transcription as a crucial effector in senescence control which is confirmed in our data showing a steep decline of *RRM2* expression levels at 48 h p.i. (p<0,0001, ANOVA) (Fig. 4C, Fig. 4D). In further support of a senescence inducing gene signature, we observed downregulation of the senescence-associated biomarker *LMNB1* both at the transcriptional (p=6.51e-08, ANOVA) and (at 72h) at protein level (Fig. 4C, Fig. 4E, Fig. 4F). In addition to the gene sets that showed a consistent up- or downregulation in MYCN-ON versus MYCN-OFF cells, we also discovered signatures with bimodal distributions. In particular, two core senescence signatures (Hernandez-Segura, A. et al., 2017 and Macedo, J. C. et al., 2018) (p<0.05) and a SASP-associated signature (Macedo, J. C. et al., 2018) (p<0.01) showed a marked drop at the 24h time point in MYCN-ON cells, only to return to baseline (MYCN-OFF) levels at later time points. A similar observation was made for gene sets that become upregulated upon *CDK4/CCND1* knockdown (p<0.001) (Fig. 4A).

**Fig 4.**
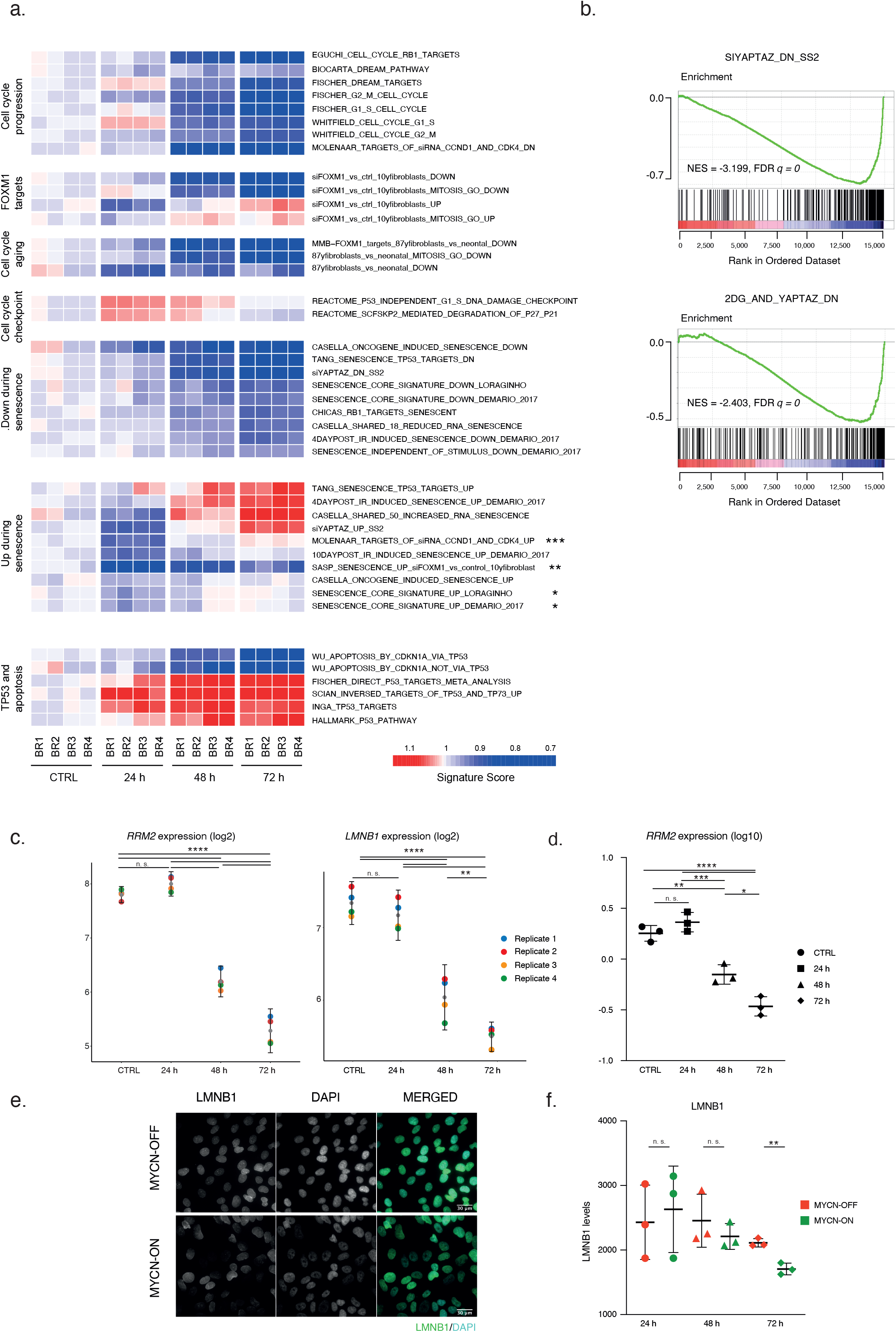
Induction of MYCN triggers a senescence-like transcriptional program and phenotype in the absence of DNA damage. (A) Heatmap representing ranksum based signature scores for cell cycle, senescence, aging, FOXM1 and p53 associated gene sets over the different timepoints p.i. MYCN activation. All depicted gene sets were significantly enriched (p<0.05), those with adjusted p-value below 0.05 are marked with an asterisk. BR= biological replicate. (B) MYCN activation correlates to inhibited dNTP metabolism as evidenced by GSEA for YAP/TAZ, which is enriched in MYCN-ON cells 72 h p.i.. (C) Log2 *RRM2* and *LMNB1* mRNA levels at 24□h, 48 h, 72 h p.i. of MYCN activation. ANOVA statistical analysis followed by a post-hoc Tukey’s test for multiple comparisons. Error bars represent 95% CI of the four biological replicates. (D) log10 *RRM2* mRNA levels 24□h, 48 h, 72 h p.i.. Results are normalised to *18s, HMBS* and *SDHA* expression. Statistical differences examined by ANOVA statistical analysis followed by a post-hoc Tukey’s test for multiple comparisons. (E) Representative images of MYCN ON and OFF cells at 72 h p.i., after immunofluorescence staining for LMNB1 (green) and counterstaining with DAPI (cyan). (F) Quantification of lamin B1 average nuclear intensity in MYCN ON and OFF cells. Statistical differences examined by unpaired Student’s t-test. Error bars represent the SD of the three biological replicates. (n.s.= not significant, * = p-value < 0, 05; ** = p-value < 0, 01; *** = p-value < 0, 001). Scale bar□=□30□μm.

To dissect the different patterns of temporal gene expression changes following MYCN activation in more detail, we applied the elbow method for k-means clustering which allows the identification of an unbiased number of clusters in a given data set (Ketchen, Jr. D. and Shook, C., 1996). Using this approach, a total of 10 clusters was identified that captured the overall variation in the bulk RNA-seq data. Four clusters (3, 6, 7 and 8) were connected to MYCN signatures (Fig. 5A, Supplementary Fig. 4A, Supplementary Table 2), while cluster 10 was strongly enriched for gene targets of cell cycle master regulators FOXM1 and E2F. Three clusters (3, 7 and 8) consisted of gene sets associated with ribosome biogenesis, a well-known MYC(N) driven process that is assumed to allow cells to cope with an increased demand for proteins required for cell growth and proliferation (Boon, K. et al., 2001) (Fig. 5A, Fig. S5A). These gene sets include *EBNA1BP2* (EBP2) (cluster 3), which stabilizes MYC and promotes MYC-mediated rRNA synthesis and cell proliferation, as well as genes encoding proteins involved in ribosome RNA transcription (*POLR3G*), processing and maturation (*WDR12, RRP1B*) and ribosome assembly and shaping (*MDN1*) (cluster 8). Genes that composed cluster 7 exhibited increased expression 24h after MYCN activation and robust downregulation at the 48 h time point. A prominent representative of this cluster was NPM1. NPM1 controls the impaired ribosome biogenesis checkpoint which becomes activated upon unscheduled MYC(N) driven ribosome production (Hald, Ø. H. et al., 2019). Checkpoint activation disables MDM2, leading to p53 stabilisation and p53 pathway activation, including p21 induction. Thus, the observed NPM1 overexpression during the very early phase of MYCN activation (24 h) suggests nucleolar stress is a driving factor in p21-driven growth delay.

**Fig 5.**
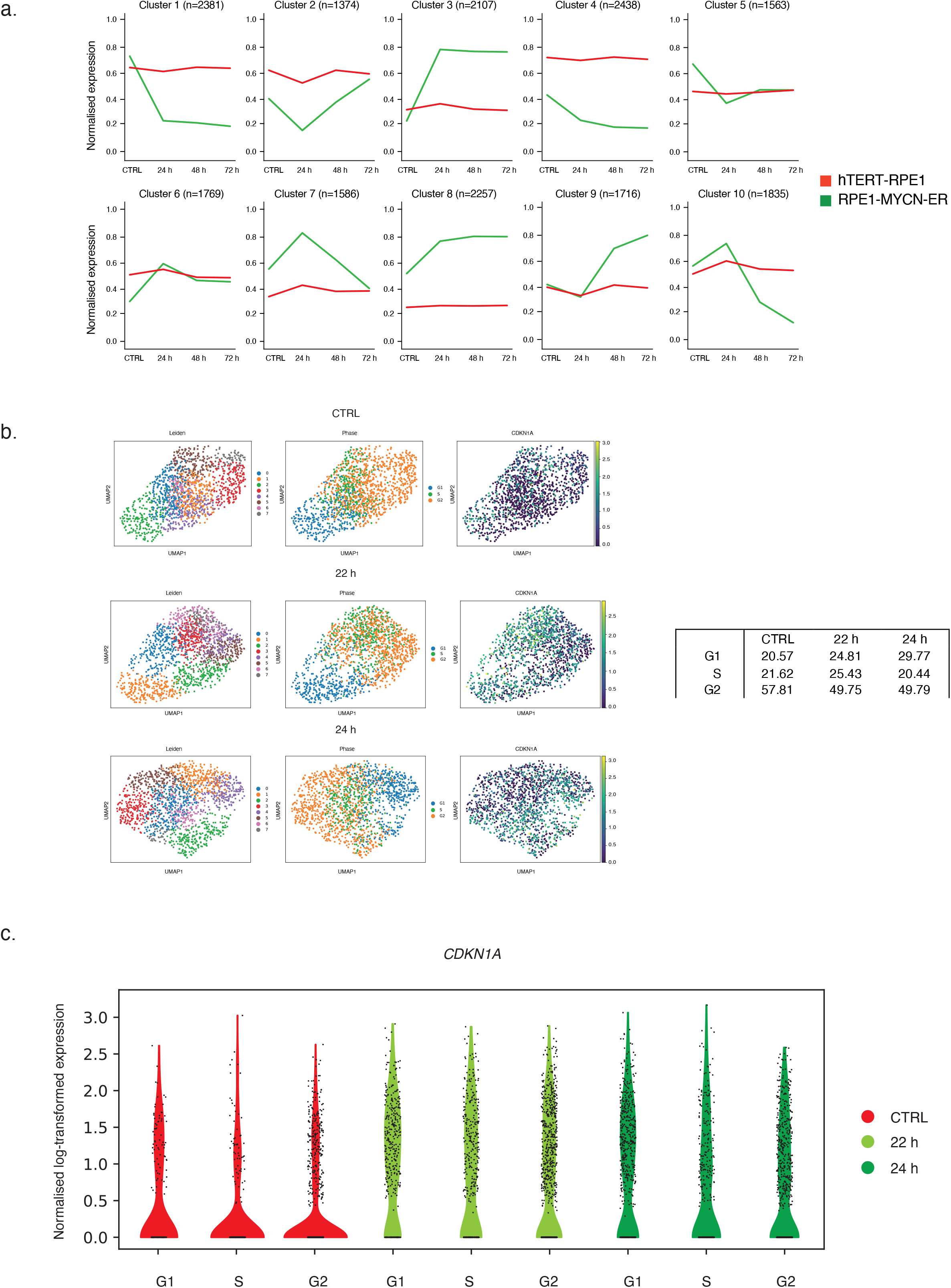
RNA-Sequencing indicates a heterogenous p21 response in MYCN-ON cells. (A) K-means clustering of all genes according to their average counts (quantile normalised) across replicates. (B) UMAP representation of leiden clusters, cell phase state and p21 expression in the four treatment conditions separately. The table below shows the cell phase distribution for the treatments. (C) Normalised log-transformed expression *CDKN1A* expression within the single cell data set for 22 and 24 h p.i..

Finally, in order to gain insight into the critical transcriptional changes and overall heterogeneity of transcriptional induction of *CDKN1A* and possible other early markers heralding the senescence transcriptome signature, we performed single cell sequencing. As *CDKN1A* was not yet significantly up 16 h p.i., but was at 24 h p.i. (Supplementary Fig. 4B), we selected two time points around *CDKN1A* inflection timepoint (22 h and 24 h). Figure 5B shows the inter-cellular heterogeneity of *CDKN1A* expression levels compared to untreated cells further supporting our immunohistochemistry staining (Fig.3D, Supplementary Fig. 3E). When correlating *CDKN1A* expression levels within cell cycle stage, the higher expression levels were proportionally increased in all stages (Fig. 5C). This indicates that *CDKN1A* expression is increasing overall, already 24 h p.i.

### 3.5. Growth attenuation in MYCN-ON cells is accompanied by cytoplasmic granules and nucleolar coalescence

The above data prompted us to further investigate known senescence hallmarks. Unexpectedly, no evidence for β-galactosidase staining was found at any of the time points following MYCN activation studied for transcriptome profiling following MYCN activation. Likewise, nuclear DAPI staining did not present evidence for senescence-associated heterochromatin foci formation (SAHFs), another canonical phenotypic senescence marker (Salama, R. et al., 2014; Swanson, E. C. et al., 2013) (Fig. S6A), and γH2AX and RPA32 staining did not reveal increased levels of DNA damage in MYCN-ON cells (Supplementary Fig. 5B-C). However, two obvious phenotypic changes were notable in MYCN-ON cells. First, we noticed a distinct nucleolar coalescence in the DAPI images of MYCN-ON cells. Quantitative immunofluorescence for fibrillarin (a protein component of C/D box small nucleolar ribonucleoproteins), confirmed this observation together with an overall increased nucleolar size (Fig. 6A-B), independently of the cell cycle status (Fig. 6C). Furthermore, a second marked change observed by transmission microscopy was a conspicuous cytoplasmic granularity in MYCN-ON cells (Fig. 6D). We reasoned that this might reflect a change in protein turnover as a consequence of the upregulated ribosomal biogenesis and nucleolar stress as also suggested by the strong NPM1 induction. However, neither staining for lysosomes, stress granules or P-bodies (Gorgoulis, V. et al., 2019; Anderson, P. et al., 2015) revealed colocalization with these granules, suggesting they do not represent typical relics of aberrant protein accrual and leaving their nature as yet undetermined (Supplementary Fig. 5D-G).

**Fig 6.**
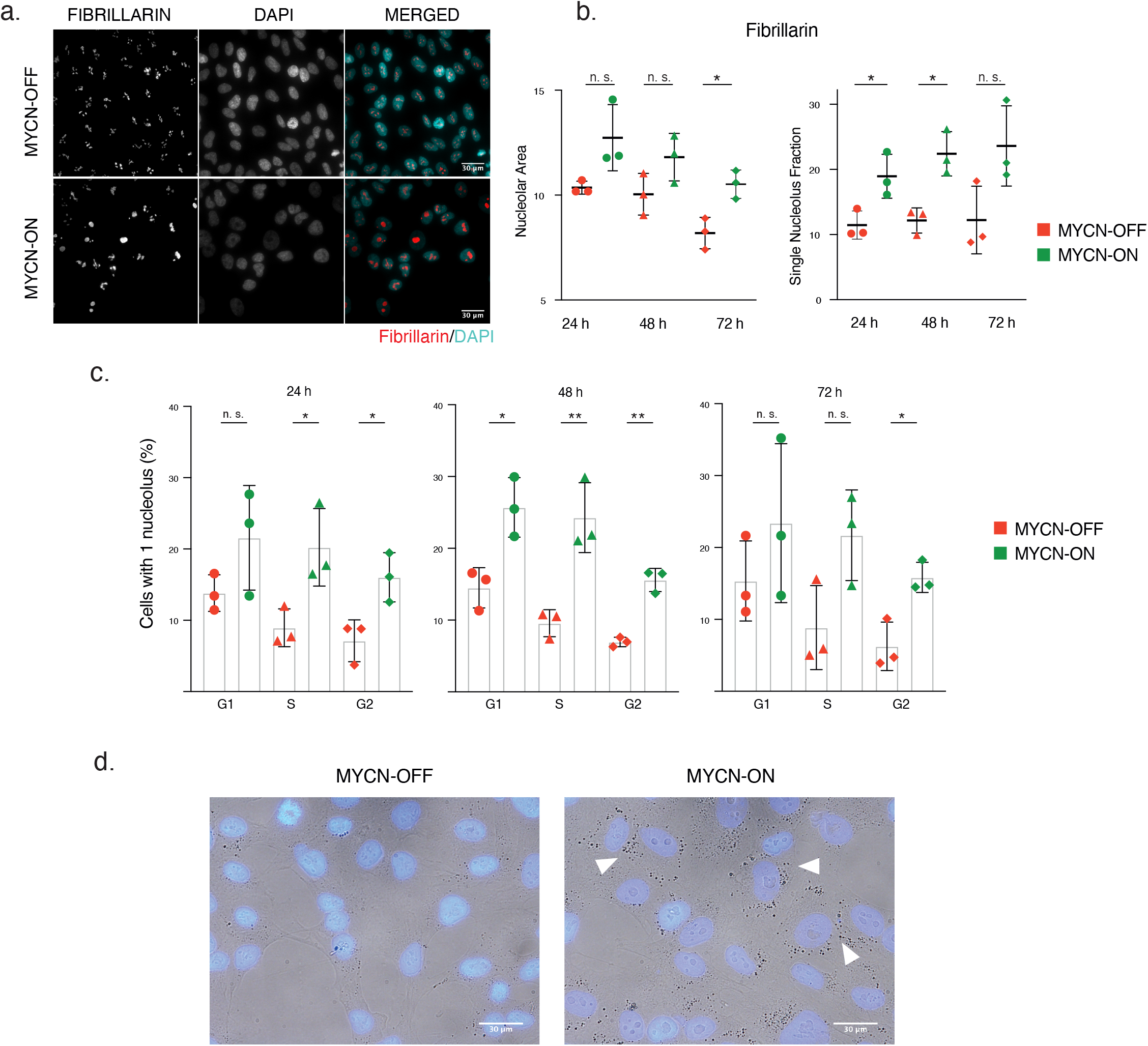
Nucleolar coalescence in MYCN-ON cells. (A) Representative images of MYCN-ON and OFF cells 72 p.i., after immunofluorescence staining for fibrillarin (red) (a protein component of C/D box small nucleolar ribonucleoproteins) counterstaining with DAPI (cyan) (B) Area of fibrillarin foci (left) and percentage of nuclei with 1 nucleolus (right) in MYCN ON versus OFF, as measured by immunofluorescence staining and quantitative image analysis. (C) DAPI-based cell cycle staging of cells with 1 single nucleolus. (D) Representative images of DAPI stained MYCN-ON and OFF cells show increase in nucleolar size and enhanced cytoplasmic granularity 72 h p.i.. Statistical differences examined by unpaired Student’s t-test. Error bars represent the SD of the three biological replicates. (n.s.= not significant, * = p-value < 0, 05; ** = p-value < 0, 01; *** = p-value < 0, 001). Scale bar□=□30□μm.

Taken together, we propose that unscheduled *MYCN* activation causes ribosome biogenesis and nucleolar stress in G2 phase, pre-priming cells for faltering in subsequent G1 and slowing down cell proliferation.

## 4. Discussion

Normal cells have tightly regulated signalling pathways that protect against insults such as unscheduled oncogene induction. Understanding how major cancer drivers such as MYCN activate and bypass such pathways during tumor initiation offers insights that may be of use for development of novel therapeutic strategies. Hence, we explored the use of immortalised neural crest-derived retinal pigment cells to investigate the early effects of MYCN activation. Unexpectedly, following the induction of MYCN, we observed attenuated cell growth without notable increased apoptosis but with a pronounced induction of p53 and p21. Attenuated proliferation was further accompanied by an increased number of cells in G1-phase and the induction of several previously reported senescence-induced gene signatures. While we observed a robust downregulation of *LMNB1*, other hallmarks of senescence (such as beta-galactosidase activity and SAHFs) were not present, suggesting that cells were not undergoing full senescence. On the other hand, MYCN-ON cells showed distinct phenotypic changes including nucleolar coalescence and cytoplasmic granularity, which aligned with transcriptional evidence for upregulation of ribosome biogenesis. Enlarged and prominent nucleoli upon increased MYCN activity have been documented before (Kobayashi, C. et al., 2015), but we now connect it to a transcriptional and phenotypic switch. We propose that ribosome biogenesis and the consequent nucleolar stress drives cells into what we propose to represent a pre-senescent state. Indeed, perturbed ribosomal biogenesis and nucleolar stress is known to drive nucleolar coalescence in senescent cells (Turi, Z. et al., 2019; Tiku, V. and Antebi, A., 2018; Chen, J. and Stark, L. A., 2019 Morlot, S. et al, 2018) and nucleolar expansion has been observed in progeria cells as a result of increased translational throughput (ribosome biogenesis and protein synthesis) (Buchwalter, A. and Hetzer, M. W., 2017). This could be mimicked by knockdown of *LMNA* and overexpression of progerin, suggesting a general influence of the lamin network on nucleolar plasticity. It is therefore tempting to speculate that the loss of lamin B1 maybe thus also act upstream of this process.

Oncogene activation (such as *MYCN*) is well known to induce alterations in ribosome biogenesis and ultimately activate the IRBC (Impaired Ribosome Biogenesis Checkpoint) as ribosomal proteins interact with the central acidic domain of MDM2 and indirectly promotes the stabilization of p53 (Turi, Z. et al., 2019). Interestingly, MYCN is implicated in upregulation of RPL (Ribosomal Protein Large) and RPS (Ribosomal Protein Small) proteins in neuroblastoma cells (Boon, K. et al.; 2001). Based on our observations, we envision that the senescence-like response triggered by MYCN activation in RPE1-MYCN-ER cells reflects a dynamic response to the oncogenic stimulus. Intriguingly, despite robust induction of *CDKN1A* in the MYCN-ON cells, neither apoptosis nor complete growth arrest was observed, and cells continued to grow, albeit more slowly. In this context, the recent discovery of a so-called “Goldilocks” zone for p21 levels was observed to control the proliferation-senescence cell-fate decision after drug treatment. Either a delayed or acute drug-induced p21 response led to senescence, while an intermediate p21 pulse enabled sustained proliferation. The cell-cycle dependent p21 overshoot that we witnessed in MYCN-ON cells may thus reflect an attempt to initiate cell cycle arrest, which was not completely successful, either by having insufficient intensity or improper timing.

In conclusion, we have identified a transient, pre-senescent state as early response to unscheduled MYCN activation. Further studies, including live cell imaging in this model and *in vivo* single cell analysis of early emerging hyperplastic lesions in MYC(N) transgenic animal tumor models can shed further light on the dynamic changes in gene activity and cellular processes that allow these cells to cope with the stress during early malignant transformation. These insights can also provide guidance towards critical dependencies and pathways as novel drug targets for MYC(N) driven malignancies, many of which are currently still associated with poor survival.

## ACKNOWLEDGMENTS

The authors would like to thank Dr. Michael D. Hogharty from the Division of Oncology at Children’s Hospital of Philadelphia for providing the immortalized MYCN-ER-hTERT RPE1 cell line. We also acknowledge A. Eggermont, and F. De Vloed for technical assistance. We would like to thank E. De Meester and S. Decraene from NXTGNT Belgium for their expertise and assistance in the sequencing experiments of this study. This research was supported by ‘Kom op tegen Kanker’ (Stand up to Cancer), the Flemish Cancer Society (Research grant to F. Speleman); Kinderkankerfonds (the non-profit childhood cancer foundation under Belgian law, Research grant to F. Speleman); the young investigator proof-of-concept project funded by the Cancer Research Institute Ghent (CRIG); Olivia Hendrickx Research Fund vzw. The following authors B. Decaesteker (1238420N), S. Vanhauwaert (12U4718N), C. Van Neste (12N6917N), M. Verschuuren (11ZF116N) and K. Durinck (12Q8319N) are supported by an FWO grant.

## Author contributions

**Sofia Zanotti:** Conceptualization, Methodology, Validation, Formal analysis (Real-time quantitative PCR, imaging data and statistical analysis), Investigation, Writing - Original Draft, Writing - Review & Editing, Visualization, Project administration; **Suzanne Vanhauwaert:** Conceptualization, Formal analysis (Real-time quantitative PCR), Writing - Original Draft, Writing - Review & Editing, Supervision, Project administration; **Christophe Van Neste:** Conceptualization, Methodology, Software, Formal analysis (Single cell RNA-sequencing data and statistical analysis), Investigation, Writing - Review & Editing, Data Curation, Visualization, Supervision; **Volodimir Olexiouk:** Software, Formal analysis (Single cell RNA-sequencing), Writing - Review & Editing; **Jolien Van Laere:** Investigation, Writing - Review & Editing; **Marlies Verschuuren:** Software, Formal analysis (imaging data and statistical analysis), Data Curation, Visualization, Writing - Review & Editing; **Liselot M. Mus:** Investigation, Visualization, Writing - Review & Editing; **Kaat Durinck:** Supervision, Writing - Review & Editing; **Laurentijn Tilleman**: Software, Formal analysis (Bulk RNA-sequencing data), Writing - Review & Editing; **Dieter Deforce:** Methodology, Resources, Supervision of NXTGNT, Writing - Review & Editing; **Filip Van Nieuwerburgh:** Conceptualization, Methodology, Resources, Supervision of NXTGNT, Writing - Review & Editing; **Michael D. Hogarty:** Methodology, Resources, Writing - Review & Editing; **Bieke Decaesteker:** Conceptualization, Methodology, Software, Formal analysis (Bulk RNA-sequencing data and statistical analysis), Data Curation, Writing - Original Draft, Writing - Review & Editing, Visualization, Supervision; **Winnok H. De Vos:** Conceptualization, Software, Formal analysis (imaging data and statistical analysis), Resources, Writing - Original Draft, Writing - Review & Editing, Supervision, Project administration, Funding acquisition; **Frank Speleman:** Conceptualization, Resources, Writing - Original Draft, Writing - Review & Editing, Supervision, Project administration, Funding acquisition.

## Abbreviations

18s: 18S ribosomal RNA
3-AP: 3-aminopyridine-2-carboxaldehyde thiosemicarbazone
4-OHT: 4-Hydroxytamoxifen
ANOVA: analysis of variance
BR: biological replicate
BSA: bovine serum albumin
CDKNA1: cyclin dependent kinase inhibitor 1A
cDNA: complementary DNA
DAPI: 4′,6-diamidino-2-phenylindole
DDX6: DEAD-box helicase 6
dH2O: distilled water
DKC1: dyskerin pseudouridine synthase 1
DMEM: Dulbecco’s Modified Eagle Medium
DMSO: dimethyl sulfoxide
DNA: Deoxyribonucleic acid
E2F1: E2F transcription factor 1
E2F2: E2F transcription factor 2
EBP2: EBNA1 binding protein 2
EDTA: Ethylenediaminetetraacetic acid
EdU: 5-ethynyl-2’-deoxyuridine
EGTA: egtazic acid
EtOH: ethanol
FBS: fetal bovine serum
FOXM1: forkhead box M1
G3BP1: G3BP stress granule assembly factor 1
GSEA: gene set enrichment analysis
h: hours
HDAC: Histone deacetylases
Hepes: 4-(2-hydroxyethyl)-1-piperazineethanesulfonic acid
HMBS: hydroxymethylbilane synthase
HRP: horseradish peroxidase
IRBC: impaired ribosome biogenesis checkpoint
KCl: potassium chloride
Ki-67: marker of proliferation Ki-67
Lamin B1: laminin subunit beta-1
MDM2: mouse double minute 2 homolog
MDN1: midasin AAA ATPase 1
mRNA: messanger RNA
mTOR: mechanistic target of rapamycin kinase
MYCN: MYCN proto-oncogene, BHLH transcription factor
NaAsO_2_: sodium arsenite
Nacl: sodium chloride
NaN_3_: sodium azide
NIH3T3: focus-forming assay
NPM1: nucleophosmin
ODC1: ornithine decarboxylase 1
OIS: oncogene-induced senescence
p21: cyclin dependent kinase inhibitor 1A
p53: tumor protein P53
PAGE: polyacrylamide gel electrophoresis
PBS: Phosphate buffered saline
PC: principal component
PCA: principal component analysis
p.i.: post induction
POLR3G: RNA polymerase III subunit G
PTMA: prothymosin alpha
qRT-PCR: quantitative real-time PCR
RAS: inducible RAS overexpression
RNA-Seq: RNA-sequencing
RNA: Ribonucleic acid
RPA32/RPA2: replication protein A 32 KDa subunit
RPE1: retinal pigmented epithelial
RPL: ribosomal protein large
RPS: ribosomal protein small
RRM2: ribonucleotide reductase regulatory subunit M2
RRP1B: ribosomal RNA processing 1B
SAHFs: senescence-associated heterochromatin foci
SASP: senescence-associated secretory phenotype
SDHA: succinate dehydrogenase complex flavoprotein subunit A T
AZ: tafazzin
TBST: Tris-buffered saline with Tween
TP53: tumor protein p53
UMAP: uniform manifold approximation and projection
WDR12: WD repeat domain 12
WT: wild type
YAP: yes associated protein 1
β-Gal: senescence-associated β-galactosidase
γH2AX: H2A histone family member X phosphorylated on Ser-14

## Supplementary Figure Legends

Supplementary Fig. 1. (A) log10 *ODC1* mRNA levels 24□h, 48 h, 72 h p.i.. Results are normalised to *18s, HMBS* and *SDHA* expression. Statistical differences examined by ANOVA statistical analysis followed by a post-hoc Tukey’s test for multiple comparisons for 24 h, 48 h, 72 h. Error bars represent the SD of the three biological replicates. (B) Log2 *MDM2, WDR12, NPM1, PTMA, DKC1* mRNA levels at 24□h, 48 h, 72 h p.i.. ANOVA statistical analysis followed by a post-hoc Tukey’s test for multiple comparisons. Error bars represent 95% CI of the four biological replicates (C) Cumulative distribution plot of the absolute log fold changes of shMYCN up-and downregulated genes in IMR-5/75 cells, shMYCN downregulated genes in SK-N-BE(2c) cells and MYCN targets in IMR-32 cells for the 24 h vs CTRL, 48 h vs CTRL and 72 h vs CTRL comparisons in RPE-1-MYC-ER cells. log2 *MYCN* mRNA levels in MYCN-ON cells. Error bars represent the 95% CI of the four biological replicates. (n.s.= not significant, * = p-value < 0, 05; ** = p-value < 0, 01; *** = p-value < 0, 001).

Supplementary Fig. 2. (A) Caspase-Glo 3/7 luminescent assay on MYCN-ON and OFF cells per time points indicated. Error bars represent the SD of the three biological replicates. (n.s.= not significant).

Supplementary Fig. 3. log2 *FOXM1, E2F1* (A), *CDKN1A* (B) and *TP53* (C) mRNA levels in MYCN-ON cells. ANOVA statistical analysis followed by a post-hoc Tukey’s test for multiple comparisons. Error bars represent the 95% CI of the four biological replicates (n.s.= not significant, * = p-value < 0, 05; ** = p-value < 0, 01; *** = p-value < 0, 001). (D) Individual quantifications of p21 (left) and p53 (right) per cell cycle phase shows that p21 is overexpressed in G1 whereas p53 is up in all phases in MYCN-ON cells. (E) Scatterplot of p21 and p53 protein expression quantification shows accumulation for both proteins in MYCN-ON cells for all time points. (n.s.= not significant, * = p-value < 0, 05; ** = p-value < 0, 01; *** = p-value < 0, 001).

Supplementary Fig. 4. (A) Gene Set Enrichment analysis on cluster 3,7 and 10 revealed enrichment for MYCN signatures and gene sets associated with ribosome biogenesis in cluster 3 and 7, while cluster 10 is enriched for FOXM1, DREAM, and E2F targets related to cell cycle. (B) log10 *CDKN1A* mRNA levels 16□h and 24 h p.i.. Results are normalised to *18s, HMBS* and *SDHA* expression. Error bars represent the SD of the two biological replicates.

Supplementary Fig. 5. (A) Representative images of DAPI channel indicating no induction of DAPI-dense nuclear foci. (B) Immunofluorescence for γH2AX (green) and RPA32 (red) foci show MYCN-ON cells do not accumulate DNA damage. (C) Quantification of γH2AX (top) and RPA32 (bottom), expressed by the number of foci per nucleus. (D-E) Imaging analysis of lysosomes show no significant increase between MYCN-ON and MYCN-OFF cells. (F) immunofluorescence staining for G3BP1 (marker for Stress Granules) (top; green) and DDX6 (marker for P-bodies) (bottom; green) merged with inverted brightfield (red) and DAPI (cyan). (G) Imaging analysis of stress granules (top) and P-bodies (bottom). Error bars represent the SD of the three biological replicates. Statistical differences examined by unpaired Student’s t-test. (n.s.= not significant, * = p-value < 0, 05; ** = p-value < 0, 01; *** = p-value < 0, 001). Scale bar□=□30□μm.

## References

Anderson, P. et al. Stress granules, P-bodies and cancer. Biochim Biophys Acta. 1849 (7), 861–870 (2015).

Baluapuri, A. et al. Target gene-independent functions of MYC oncoproteins. Nat Rev Mol Cell Biol. 21, 255–267 (2020).

Boon, K. et al. N-myc enhances the expression of a large set of genes functioning in ribosome biogenesis and protein synthesis. EMBO J. 20 (6), 1383–1393 (2001).

Brady, S. W. et al. Pan-neuroblastoma analysis reveals age- and signature-associated driver alterations. Nat Commun. 11 (1), 5183 (2020).

Buchwalter, A., and Hetzer, M.W. Nucleolar expansion and elevated protein translation in premature aging. Nat Commun. 8 (1), 328 (2017).

Chen, J. and Stark, L. A. Insights into the Relationship between Nucleolar Stress and the NF-κB Pathway. Trends in Genetics. 35 (10), 768–780 (2019).

Cole, K. A. et al. RNAi screen of the protein kinome identifies checkpoint kinase 1 (CHK1) as a therapeutic target in neuroblastoma. Proc Natl Acad Sci U S A. 108 (8), 3336–3341 (2011).

De Vos, W. H. et al. High content image cytometry in the context of subnuclear organization. Cytometry A. 77 (1), 64–75 (2010).

Dobin, A. et al. STAR: ultrafast universal RNAseqaligner. Bioinformatics. 29 (1), 15–21 (2013).

Fredlund, E. et al. High Myc pathway activity and low stage of neuronal differentiation associate with poor outcome in neuroblastoma. PNAS. 105 (37), 14094–14099 (2008).

Georgakilas, A. G. et al. p21: A Two-Faced Genome Guardian. Trends Mol Med. 23 (4), 310–319 (2017).

Gorgoulis, V. et al. Cellular Senescence: Defining a Path Forward. Cell. 179 (4), 813–827 (2019).

Hald, Ø. H. et al. Inhibitors of ribosome biogenesis repress the growth of MYCN-amplified neuroblastoma. Oncogene. 38 (15), 2800–2813 (2019).

Hernandez-Segura, A. et al. Unmasking Transcriptional Heterogeneity in Senescent Cells. Curr Biol. 27 (17), 2652–2660 (2017).

Hiemer, S. E. et al. A YAP/TAZ-Regulated Molecular Signature Is Associated with Oral Squamous Cell Carcinoma. Mol. Cancer Res. 13 (6), 957–968 (2015).

Hovestadt, V. et al. Medulloblastomics revisited: biological and clinical insights from thousands of patients. Nat Rev Cancer. 20 (1), 42–56 (2019).

Ketchen, Jr., D. and Shook, C. The application of cluster analysis in strategic management research: an analysis and critique. Strateg. Manag. J. 17 (6), 441–458 (1996).

Kobayashi, C et al. Enlarged and prominent nucleoli may be indicative of MYCN amplification: a study of neuroblastoma (Schwannian stroma-poor), undifferentiated/poorly differentiated subtype with high mitosis-karyorrhexis index. Cancer. 103 (1), 174–80 (2005).

Li, B. and Dewey, C. N. RSEM: accurate transcript quantification from RNAseqdata with or without a reference genome. BMC Bioinformatics. 12 (1), 323 (2011).

Macedo, J.C. et al. FoxM1 repression during human aging leads to mitotic decline and aneuploidy-driven full senescence. Nat Commun. 9, 2834 (2018).

Martin, M. et al. Cutadapt removes adapter sequences from high-throughput sequencing reads. EMBnet.journal. 17 (1) (2011).

Morlot, S. et al. Nucleolar stress triggers the irreversible cell cycle slow down leading to cell death during replicative aging in Saccharomyces cerevisiae. BioRxiv. 297093 (2018).

Oliynyk, G. et al. MYCN-enhanced Oxidative and Glycolytic Metabolism Reveals Vulnerabilities for Targeting Neuroblastoma. iScience. 21, 188–204 (2019).

Pont, F. et al. Single-Cell Signature Explorer for comprehensive visualization of single cell signatures across scRNA-seq datasets. Nucleic Acids Res. 47 (21), e133 (2019).

Rajbhandari, P. et al. Cross-Cohort Analysis Identifies a TEAD4–MYCN Positive Feedback Loop as the Core Regulatory Element of High-Risk Neuroblastoma. Cancer Discov. 8 (5), 582–599 (2018).

Rickman, D. S. et al. The Expanding World of N-MYC–Driven Tumors. Cancer Discov. 8 (2), 150–163, (2018).

Roukos, V. et al. Cell cycle staging of individual cells by fluorescence microscopy. Nat Protoc. 10, 334–348 (2015).

Rufini, A. et al. Senescence and aging: the critical roles of p53. Oncogene. 32, 5129–5143 (2013).

Salama, R. et al. Cellular senescence and its effector programs. Genes Dev. 28, 99–114 (2014).

Swanson, E. C. et al. Higher-order unfolding of satellite heterochromatin is a consistent and early event in cell senescence. J Cell Biol. 203 (6), 929–42 (2013).

Tiku, V. and Antebi, A. Nucleolar function in lifespan regulation. Trends in biology. 28 (8), 662–672 (2018).

Todaro, G. J. and Green, H. Quantitative studies of the growth of mouse embryo cells in culture and their development into established lines. J. Cell. Biol. 17 (2), 299–313 (1963).

Turi, Z. et al. Impaired ribosome biogenesis: mechanisms and relevance to cancer and aging. Aging (Albany NY) 11 (8), 2512–2540 (2019).

Wang, Q. et al. ID2 Expression Is not Associated with MYCN Amplification or Expression in Human Neuroblastomas. Cancer Res. 63 (7), 1631–1635 (2003).

Westermann, F. et al. Distinct transcriptional MYCN/c-MYC activities are associated with spontaneous regression or malignant progression in neuroblastomas. Genome biology. 9 (10) (2008).

Xie, Q. et al. YAP/TEAD-mediated transcription controls cellular senescence. Cancer Res. 73 (12), 3615–3624 (2013).

Yue, M. et al. Oncogenic MYC Activates a Feedforward Regulatory Loop Promoting Essential Amino Acid Metabolism and Tumorigenesis. Cell Rep. 21 (13), 3819–3832 (2017).

