## Supplementary figures and images for "MYCN-induced nucleolar stress drives an early senescence-like transcriptional program in hTERT-immortalized RPE cells"

### Supplementary Figure 1

Supplementary Fig. 1

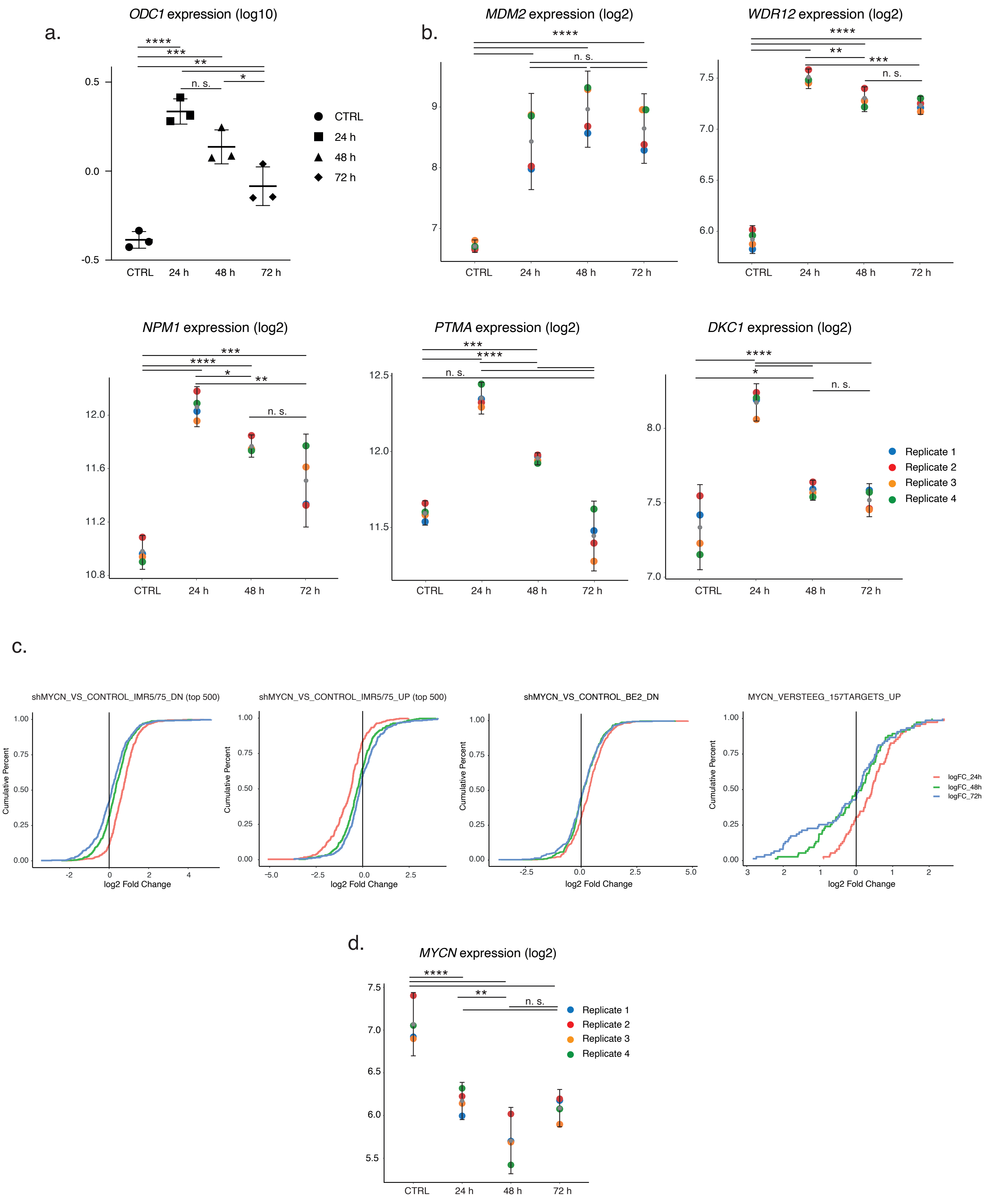

### Supplementary Figure 2

Supplementary Fig. 2

a.

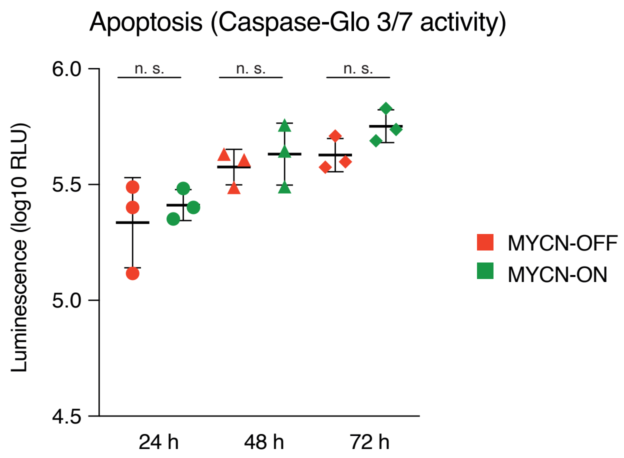

### Supplementary Figure 3

#### Supplementary Fig. 3

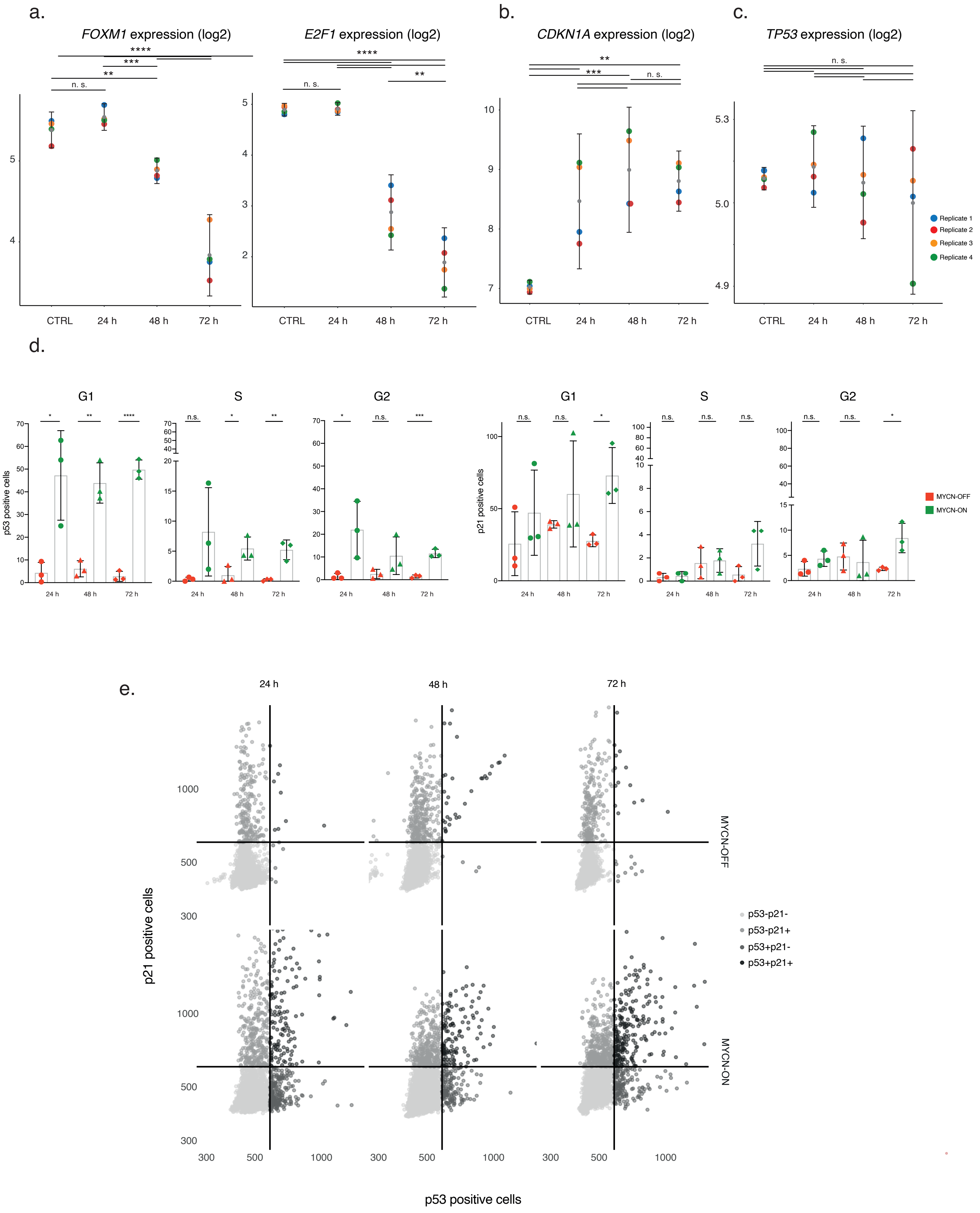

### Supplementary Figure 4

#### Supplementary Fig. 4

a.

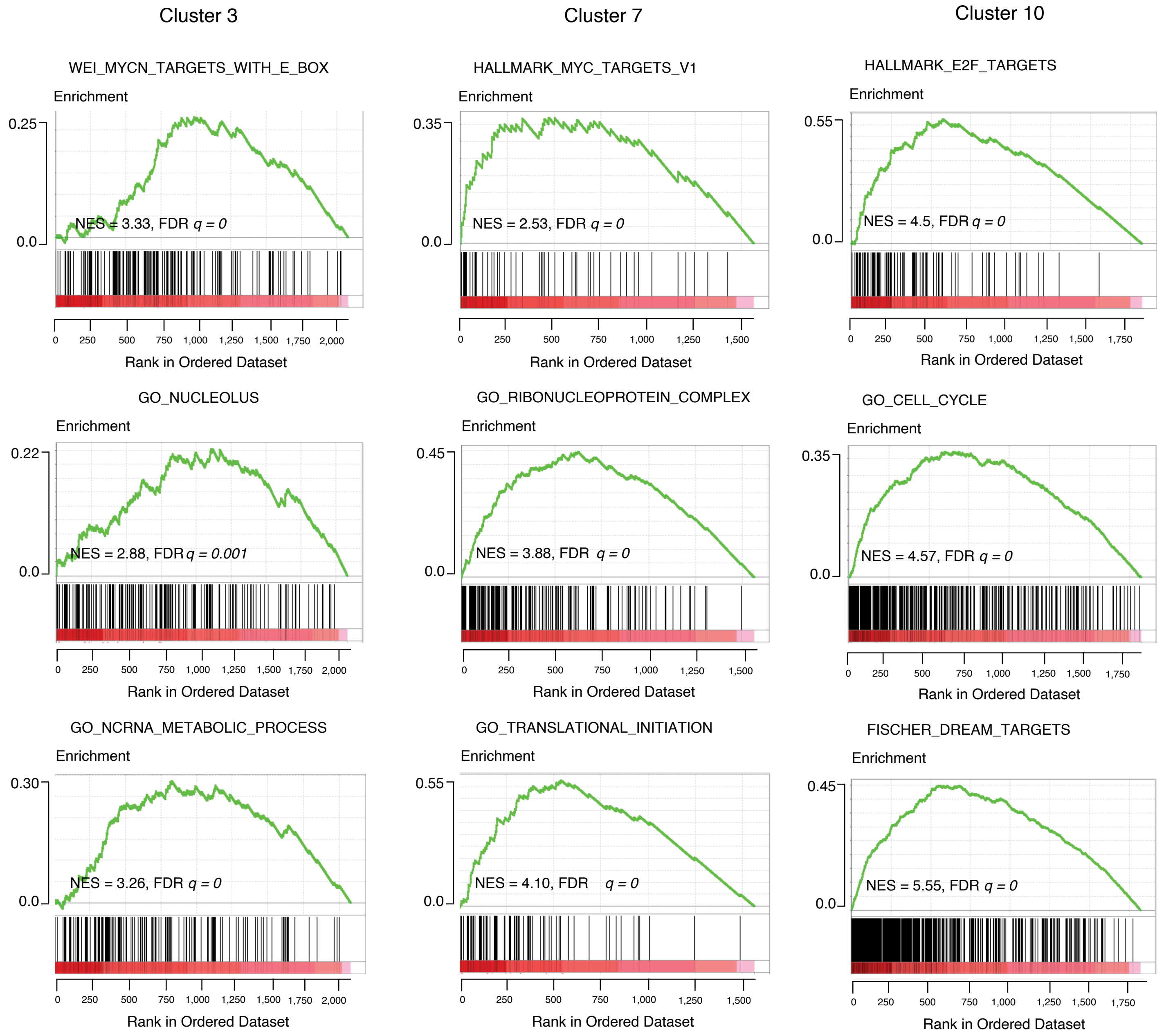


b.

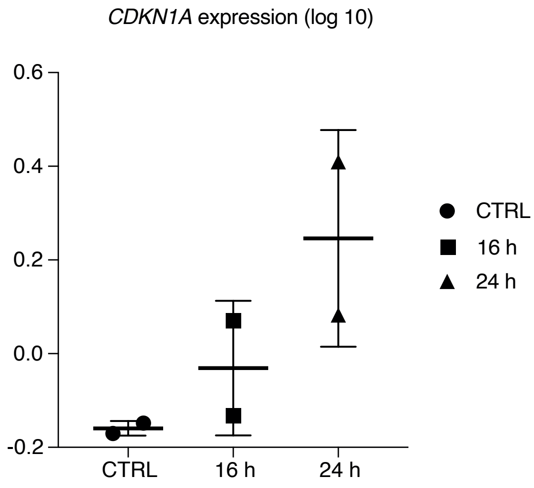

### Supplementary Figure 5

Supplementary Fig. 5

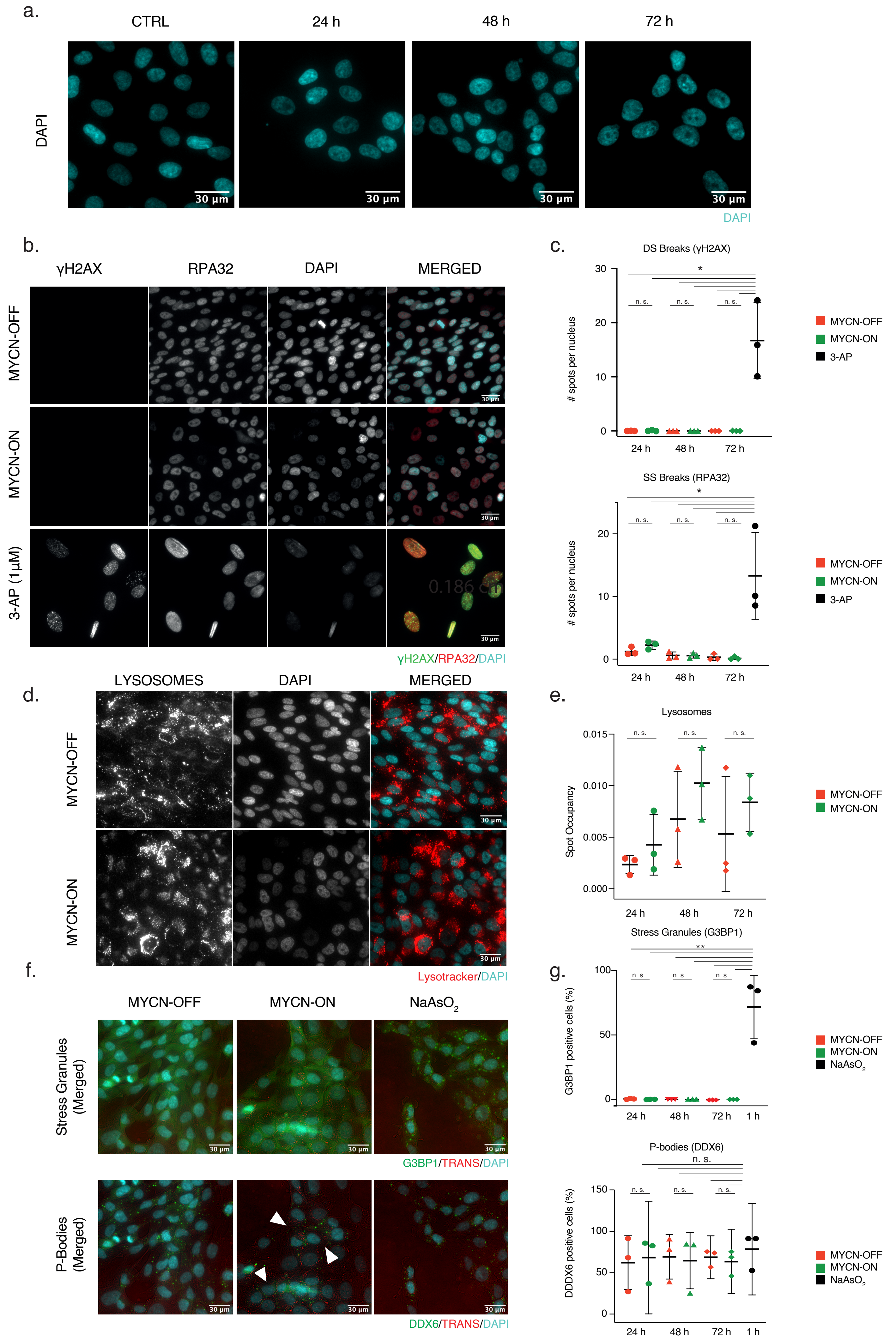
